# Musical training-induced plasticity reflects melodic rather than instrumental expertise

**DOI:** 10.64898/2026.07.29.741416

**Authors:** Tamar Ben David, Shachar Gal, Romi Kaplan, Danielle Catalogna, Asaf Madar, Niv Tik, Michal Bernstein-Eliav, Ido Tavor

## Abstract

Musical training produces behavioral gains that extend beyond the practiced context. Yet, it is unclear whether such generalization is accompanied by broadly transferable neural changes or whether training-induced plasticity remains tied to the specific features of the learned material. In this study, 80 musically naïve adults completed four piano-training sessions and underwent functional MRI before and after training. Twenty untrained participants served as passive controls. Participants performed a passive listening task in which they listened to the trained melody (Für Elise) and an untrained melody (Ode to Joy), each played on a trained musical instrument (piano) and an untrained musical instrument (saxophone). Following training, participants demonstrated improved accuracy and rhythmic performance when playing an unfamiliar melody, indicating behavioral generalization. Training-induced neural changes, however, were predominantly associated with the learned melodic content. Listening to Für Elise elicited increased activity in the intraparietal and supplementary motor cortex and in the cerebellum, as well as decreased activity in the superior temporal cortex. Consistently, pre- and post-training scans were most accurately classified based on brain activity to Für Elise played on piano or saxophone, rather than brain response to Ode to Joy, suggesting that neural reorganization was driven more strongly by the practiced melody than by the trained instrumental timbre. Training also enhanced the neural separability of both melodic and instrumental information. Critically, pre-training responses to Für Elise played on the piano predicted subsequent playing accuracy, whereas pre-training sensitivity to instrumental timbre predicted rhythmic performance in a novel melodic context. Together, these findings suggest that while acquired musical skills behaviorally transfer to novel material, practice-induced functional plasticity remains predominantly melody-specific. We conclude that musical skill acquisition reflects complementary contributions of pre-existing neural characteristics and learning-induced plasticity, with baseline auditory representations potentially supporting successful behavioral transfer.

## Introduction

Experience-dependent neuroplasticity is a core mechanism supporting human learning. Over the past couple of decades, functional reorganization following learning has been extensively explored using functional MRI (fMRI). FMRI studies have provided evidence for learning-induced changes in brain activity in skill-related regions across a variety of cognitive domains (Tardif et al., 2016), including language learning (Alotaibi et al., 2023; Coldham et al., 2025), motor skill acquisition (Dayan and Cohen, 2011; Ossmy and Mukamel, 2018, 2016), mathematical problem solving (Emerson and Cantlon, 2015; Kuhl et al., 2020; Rivera et al., 2005) and programming (Hishikawa et al., 2023). A particularly widely used framework for studying the neural mechanisms underlying learning is musical training, as playing a musical instrument requires dynamic interactions among motor coordination, auditory processing, spatial perception, cognitive control, and memory (Herholz and Zatorre, 2012; Olszewska et al., 2021). Consequently, investigating the neural modifications associated with acquiring musical skills may uncover broader principles of experience-dependent plasticity.

Evidence from cross-sectional and longitudinal studies provides complementary perspectives on the neural basis of musical expertise. Cross-sectional studies comparing expert musicians to musically naïve individuals have consistently shown enhanced brain activity in skill-specific regions, including auditory (e.g., superior temporal gyrus, planum temporale), motor (primary and supplementary motor cortices), and parietal areas involved in spatial and sensorimotor processing during passive music listening (Bianchi et al., 2017; Criscuolo et al., 2022; Olszewska et al., 2021; Pando-Naude et al., 2021; Vuust et al., 2022). Longitudinal studies, which have followed musically naïve participants over time to separate the effects of training from pre-existing neural characteristics, have revealed significant cortical (e.g., auditory and motor areas) and subcortical (e.g., cerebellar) adaptations even after short-term piano learning interventions. These included both increased activity (Brown and Penhune, 2018; Li et al., 2019, 2018; Olszewska et al., 2021; Zatorre et al., 2007) and, in some cases, decreased responses interpreted as improved neural efficiency (Chen et al., 2012). The above studies have primarily focused on passive listening to a trained melody. Therefore, it remains unclear whether the observed effects are specific to the practiced melody, specific to the practiced musical instrument, or may generalize beyond the training context. An important aspect of skill acquisition is the degree to which learning transfers to novel settings, including unlearned melodies and instruments. Generalization of musical training has been observed at the behavioral level, such that learners can perform unlearned melodies or generalize across listening conditions (Rohrmeier and Rebuschat, 2012; Van Hedger et al., 2023), and at the neural level, where training was shown to reshape the neural representation of untrained melodies (Freitas et al., 2018). However, previous studies have typically utilized untrained melodies merely as a control condition with low or no significant effect in comparison to the trained melody results (Herholz et al., 2016; Olszewska et al., 2021; Wollman et al., 2018).

To answer the question of specificity versus generalization in musical training, the present study tests changes in brain activity in response to learned and unlearned melodic and instrumental contexts. We further investigate the predictive value of the response to untrained melodies and instruments for subsequent performance in learned and unlearned contexts. By that, we examine whether there is a neural predisposition to learning music such that brain activity patterns before learning can predict future success. While longitudinal studies typically use pre-learning activity only as a baseline assessing training-related change, we propose that individual differences in pre-training neural responses may also carry meaningful information on subsequent learning outcomes. Therefore, analyzing individual differences at baseline may provide insights into variations in learning trajectories and enable predictive modeling.

## Methods

### Participants

Eighty young healthy adults were recruited for the piano training group (mean age: 26.025; SD: 3.811; 40 females, all right-handed). An additional twenty participants were recruited for the control group (mean age: 25.885; SD: 3.197; 12 females, all right-handed). All participants had no previous musical experience and no history of neurological disease or psychological disorders, drug or alcohol abuse, or use of neuropsychiatric medication. The research protocol was approved by the Institutional Review Board of the Sheba Medical Center (Ramat Gan, Israel), and all participants provided written informed consent prior to their participation. Participants were compensated at a rate of 50 NIS per hour, totaling 350 NIS for the training participants (for their involvement in four training sessions and two MRI sessions) or 150 NIS for the control participants (for two MRI sessions).

### Experimental design

Participants of the piano training group attended four sessions (mean number of days between first and last session: 12, SD: 3). All meetings included a piano training session, while the first and last meetings also included MRI scans (Figure 1A).

**Figure 1.**
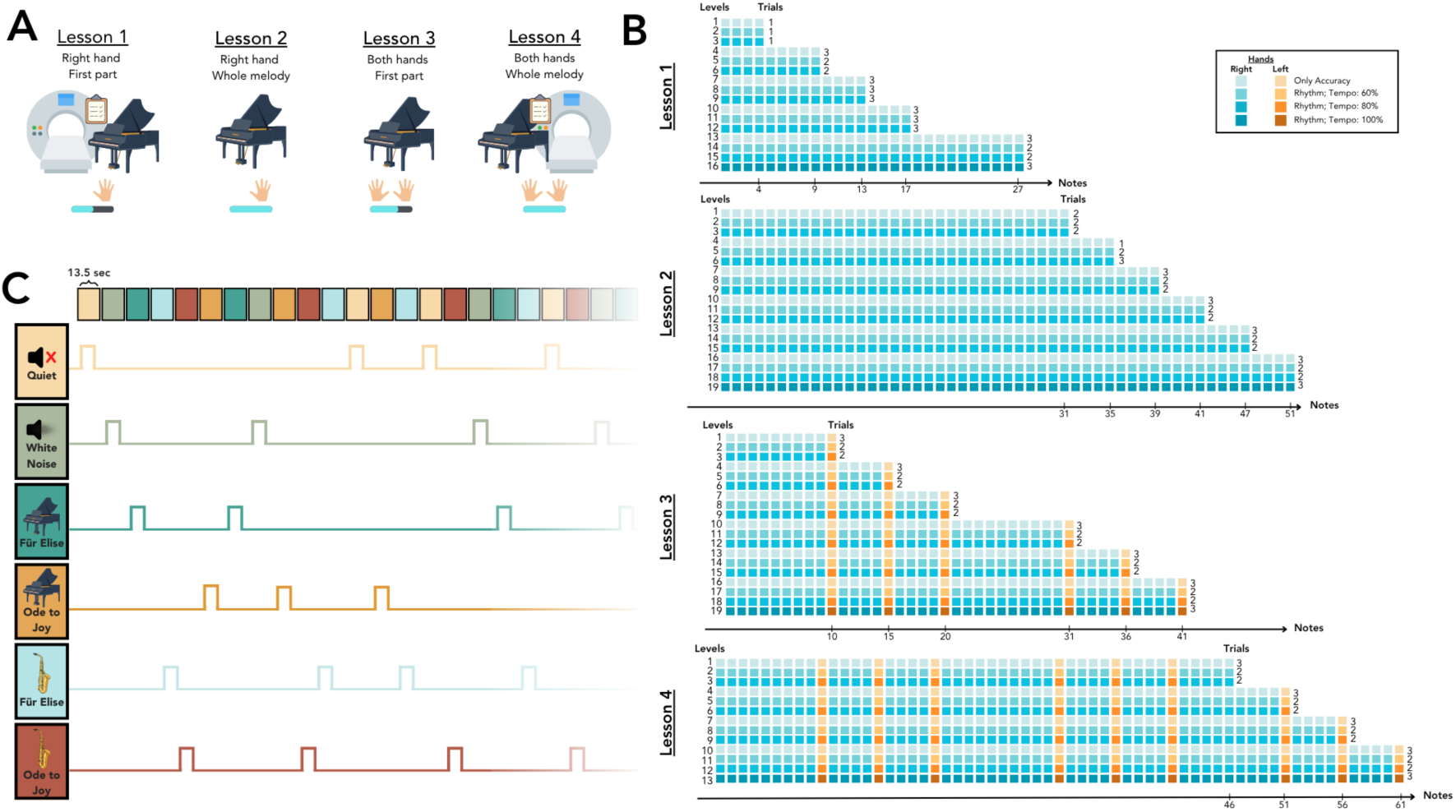
Experimental design and learning procedure. **A.** Participants in the piano training group attended four sessions. The first session began with a pre-training MRI scan followed by a pre-training test, in which participants played an unfamiliar original melody composed by the research team. They then learned to play the first half of *Für Elise* using their right hand only. The second session included piano training only, and at its end participants played the entire musical piece using their right hand (51 notes). In the third session, participants learned to play the first half of the musical piece using both hands simultaneously, and in the fourth session they played the entire piece using both hands (61 notes). They then performed the post-training test and a post-training MRI scan. **B.** The fMRI task included six auditory stimuli: *Für Elise* played on the piano or saxophone, *Ode to Joy* played on the piano or saxophone, and quiet and white noise, which were presented in a blocked design. **C.** Für Elise learning progression. Each row corresponds to a different level. Within each row, the bars represent individual notes, and the total horizontal length reflects how many notes are included in that level. The number shown at the far right indicates the number of trials for that level. The color shading of each row depicts the rhythm and tempo requirements. Blue bars indicate notes played with the right hand, while orange bars indicate notes played with the left hand.

### Piano training procedure

Participants underwent a training protocol consisting of four 75-minute lessons using ‘*Pyano*’, an in-house program designed to teach the first 61 notes of Ludwig van Beethoven’s *Für Elise* in a gradual, uniform manner (Figure 2). The protocol progressed from right-hand performance (first 51 notes; Lessons 1–2) to bimanual playing (61 notes; Lessons 3–4), with each lesson divided into 13–19 levels containing 1–3 identical trials. Each level commenced with a video tutorial demonstrating the required tempo and fingering, followed by an animation highlighting the notes on a virtual keyboard to guide placement and sequence. Difficulty increased between levels through either the addition of new notes or an increase in tempo (Figure 1C). Performance was monitored via real-time feedback where note accuracy errors were highlighted in red and rhythmic errors in blue. Progression to the next trial or level required achieving perfect note accuracy, with erroneous trials repeated until corrected, but did not require the absence of rhythmic errors.

**Figure 2.**
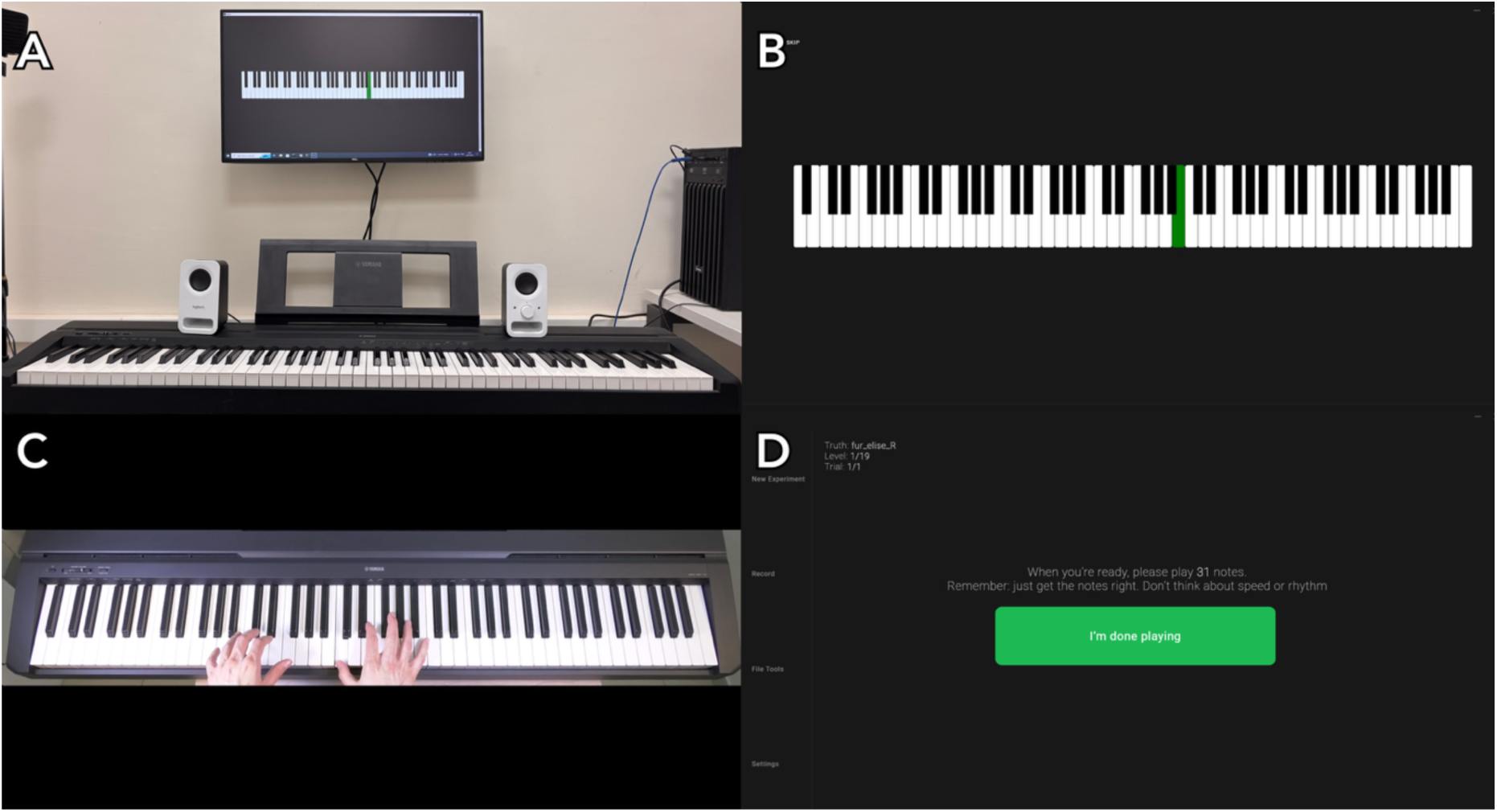
*Pyano* setup and examples. **A.** Piano training setup, including a computer running the *Pyano* training software, a screen and two speakers. A Yamaha P-45 electric piano was connected to the computer via Bluetooth. **B.** Screenshot of the video tutorial in *Pyano*. **C.** Screenshot of the animation tutorial in *Pyano.* **D.** Screenshot of the instructions presented to participants before playing the required notes.

### Performance generalization tests

To assess baseline piano proficiency and the ability to generalize skills beyond the trained musical piece, participants completed two generalization tests using the *Pyano* software: one administered before the initial training session and the other after the final session. These tests utilized two distinct 16-note melodies composed by the research team, which were counterbalanced across participants. Each test consisted of six trials structured by increasing sequence length: two trials of 9 notes, two of 13 notes, and two of 16 notes. Unlike the training protocol described above, these tests did not provide corrective feedback or allow for the repetition of unsuccessful trials; instead, participants were limited to six fixed attempts, with the target sequence replayed before each attempt.

### Behavioral data analysis

Three behavioral measures were calculated to evaluate learning progress and compare pre- and post-learning performance:

***Accuracy score*** was calculated based on Levenshtein’s ratio (Levenshtein, 1966). Sequences of notes played by participants were analyzed to detect the longest series matching the original composition, to minimize errors from omissions or additions. If an error was detected, the next longest matching series was searched for, and so on. This iterative process identified the most accurate sequence performed in each trial. An accuracy score, ranging from 0 to 1, was calculated by dividing the number of matching notes by the total notes to be played. Accuracy was calculated separately for each trial, level, and lesson.

***Rhythm score*** was computed by first measuring the time intervals between consecutive notes and normalizing them to the played tempo. These adjusted intervals were then compared to the expected gaps in the original melody, and the deviation between the two was calculated. To obtain the final rhythm score, we averaged the deviations across each trial, level, and lesson. To facilitate a more intuitive interpretation, whereby higher values reflect better rhythmic accuracy, the deviation was transformed into its complement, yielding the final rhythm matching score, ranging from 0 to 1.

***Tempo score*** was calculated by measuring the timing of the correctly played sequence from start to end. A tempo score was calculated across each trial, level, and lesson.

Paired t-tests were used to compare pre- and post-learning performance (i.e., accuracy, rhythm, and tempo scores) across participants.

### MRI acquisition

MRI scans were performed at the Tel Aviv University Strauss Center for Computational Neuroimaging using a 3T Magnetom Siemens Prisma scanner (Siemens, Erlangen, Germany) with a 64-channel RF coil. Participants underwent two MRI sessions, before and immediately after the piano training procedure (see Figure 1). The MRI protocol included T1-weighted, T2- weighted, and functional MRI, as well as Diffusion Weighted Imaging (DWI) scans not analyzed in the current work.

T1-weighted images were acquired using a 3D magnetization-prepared rapid acquisition gradient echo (MPRAGE) sequence, with the following parameters: voxel size. T2-weighted images were acquired using a SPACE (SPC) sequence, with the following parameters: voxel size. T2*-weighted images were acquired using a gradient-echo echo-planar protocol (GE-EPI) with the following parameters:, Multiband acceleration Factor (Simultaneous Multi-Slice) of 8, voxel size, flip angle: 52°, AP phase encoding. A field map was acquired using spin-echo EPI images with opposite phase-encoding directions.

### Functional MRI task

During fMRI scans, participants completed a passive listening task where they were presented with six types of auditory stimuli via MRI-compatible earphones: quiet, white noise, *Für Elise* played on the piano or saxophone, and *Ode to Joy* played on the piano or saxophone (Figure 1B). All participants were familiar with the two melodies prior to the experiment, as verified by familiarity ranking higher than 4 on a 1-5 scale, ensuring comparable baseline familiarity. Each stimulus lasted 13.5 seconds and was repeated seven times in a counterbalanced order across subjects. Participants were instructed to focus on a cross displayed in the center of the screen, listen attentively to the auditory stimuli, and press a button whenever the stimulus changed to ensure attentiveness.

### MRI data preprocessing

All data were preprocessed using the Human Connectome Project (HCP) minimal preprocessing pipeline, which is based on FSL and FreeSurfer functions (Glasser et al., 2013). The functional data preprocessing included motion and distortion corrections, nonlinear alignment to MNI space, and denoising with FMRIB’s ICA-based Xnoiseifier (FIX) (Griffanti et al., 2014; Salimi-Khorshidi et al., 2014). The data were then projected onto a surface representation of 91,282 vertices (“grayordinates”) in standard space, followed by alignment and registration using Multimodal Surface Matching (MSMAll) (Robinson et al., 2018, 2014).

### MRI data analysis

The task-fMRI data were analyzed as a block design using the HCP’s pipeline, with activity estimates computed using a general linear model (GLM) in FSL’s FILM (Jenkinson et al., 2012; Woolrich et al., 2009). Stimuli were convolved with a double gamma hemodynamic response function (Glover, 1999) to generate the main model regressors. Task-contrasts comparing each auditory condition with the quiet baseline condition (e.g., *Für Elise* > Quiet; *Ode to Joy* > Quiet; Piano > Quiet; Saxophone > Quiet) were constructed to examine the neural patterns evoked by each stimulus independently. In addition, auditory conditions were directly contrasted (e.g., *Für Elise* > *Ode to Joy*; Piano > Saxophone) to probe changes in neural representations across different musical contexts.

A vertex-wise GLM was used to compare pre- and post-learning whole-brain activity maps for the following task contrasts: *Für Elise* piano > quiet; *Für Elise* > *Ode to Joy*; *Für Elise* piano > *Ode to Joy* piano; piano > saxophone; *Für Elise* piano > *Für Elise* saxophone. These task contrasts were selected to reflect learning-induced activity changes in response to either the learned melody, the learned musical instrument, or both. Statistical significance was assessed using vertexwise two-tailed paired t-tests, comparing pre- and post-training activations. Vertices showing a significant change in activity were identified at a threshold of, corrected for multiple comparisons using the False Discovery Rate (FDR) correction.

### Activity-based classification of pre- and post-training scans

We tested whether scans acquired before and after training could be differentiated based on whole-brain activity patterns in response to the following task conditions: *Für Elise* played on the piano, *Für Elise* played on the saxophone, *Ode to Joy* played on the piano, and *Ode to Joy* played on the saxophone, each contrasted against quiet. Separate models were trained for each task condition, as well as a combined model incorporating all four. Models were trained in a repeated 10-fold cross-validation (CV) routine (10 folds × 10 repetitions, following Lumumba et al., 2024) to classify scans as pre- or post-training. First, principal component analysis (PCA) was applied to the training set to reduce data dimensionality in the task- contrast activation maps, retaining components that explained 80% of the variance. The resulting component loadings were regressed against the individual data of the held-out test-set to compute predictive features called “expression scores”, following the brain-basis set (BBS) pipeline (Sripada et al., 2020). These features served as input for an elastic-net logistic regression classifier. Across repetitions, fold assignments were reshuffled to improve robustness and reduce variance in performance estimates. Hyperparameter tuning of the regularization parameter was performed within each training fold using a nested cross- validation procedure, ensuring that the test data remained fully independent from both model fitting and parameter selection. To avoid data leakage, the two scans of each individual subject were kept in the same data fold. Classification performance was assessed using the receiver operating characteristic (ROC) area under the curve (AUC) and mean accuracy across folds; statistical significance was estimated via permutation testing (10,000 label shuffles). To determine chance-level performance, an additional classification model was trained on data from the passive control participants, in which no differences in brain activity are expected. Differences in classification performance between task-condition models were assessed using pairwise paired t-tests on the accuracy values obtained from corresponding cross-validation folds and repetitions. All possible pairs of models were compared, and the resulting p-values were corrected for multiple comparisons using the Benjamini-Hochberg false discovery rate procedure.

### Prediction of post-training behavioral performance from pre-training brain activity

We tested whether scans acquired before training could be used to predict performance accuracy scores in the final lesson, in which the entire musical piece was performed bimanually. The following task conditions were used: *Für Elise* played on the piano, *Für Elise* played on the saxophone, *Ode to Joy* played on the piano, and *Ode to Joy* played on the saxophone, each contrasted against quiet. Voxel selection was performed within each training fold based on Pearson’s correlations with the target variable. Voxels were retained based on a threshold on the absolute correlation, with the threshold treated as a hyperparameter and optimized via nested cross-validation ().

Dimensionality reduction was performed using PCA, and the resulting features were mean-centered and normalized. A linear regression model with elastic net regularization was applied. The regularization strength (**λ**) and the optimal number of features were determined through a cross-validation hyperparameter tuning process ( with 30 log-spaced points and ), ensuring a balance between model complexity and predictive performance. Prediction accuracy was measured as the Pearson’s correlation coefficients between predicted and actual behavioral scores, as well as the mean squared error (MSE).

Five models were examined in total, based on brain activity to the four task contrasts mentioned above as well as a combined model, therefore significance levels of the correlations between actual and predicted behavioral scores were corrected for multiple comparisons using the Benjamini–Hochberg false discovery rate (FDR) procedure. We then compared predictive performance across all possible pairs of models using paired t-tests, and the resulting p-values were also FDR corrected for multiple comparisons.

### Contribution maps calculation

Contribution maps, depicting brain regions where activity most strongly influenced models performance (i.e., classification of pre- vs. post-training scans, or predicting accuracy scores), were created using the PCA loadings multiplied by the classification or prediction model coefficients, averaged over all repetitions and cross-validation folds.

### Activity-based classification of melodies and musical instruments

To examine how piano training influenced the neural representations of melodies and musical instruments, we examined classification of melodic and instrumental content based on activation patterns before or after training. Two separate binary classification tasks were examined: one aimed to differentiate between two melodies (*Für Elise* vs. *Ode to Joy*), and the other to differentiate between two instruments (Piano vs. Saxophone). Each classification task was performed independently on data from the pre-training and post-training sessions. Separate classification models were constructed for each of the task conditions contrasted against quite. Classification was evaluated using 10-fold CV routine (10 folds × 10 repetitions) with the two scans of each individual kept in one fold to avoid data leakage. Within each fold, dimensionality was reduced using PCA, retaining components that accounted for 80% of the variance. The resulting component were used to generate “expression scores” (Sripada et al., 2020) which served as input for an elastic-net logistic regression classifier. Hyperparameter tuning of the regularization parameter was performed within each training fold using a nested cross-validation procedure. Model performance was quantified using the AUC-ROC and accuracy; statistical significance was evaluated using permutation testing (10,000 permutations). Additional classification models were trained on data from the passive control participants only, serving as baseline (chance-level) performance. Classification performance pre- vs. post-training were compared using paired t-tests. Voxel-wise contribution weights were computed by projecting regression coefficients back to the original feature space. These contribution maps were averaged across CV iterations and visualized on the cortical surface. All analyses were implemented in MATLAB.

## Results

### Behavioral performance

Following four piano training sessions, all participants demonstrated improvement in performance of the learned melody, *Für Elise,* and were able to play it to varying degrees of proficiency. Specifically, 65 of the 80 participants successfully completed the first two lessons using only their right hand (Figure 3A, top). After completing all four sessions and performing the melody in the final two lessons with the addition of the left hand, 68 participants were able to play the piece bimanually (Figure 3A, bottom). The remaining participants nevertheless learned to play the melody but did not achieve an error-free performance within the time allocated for each lesson.

**Figure 3.**
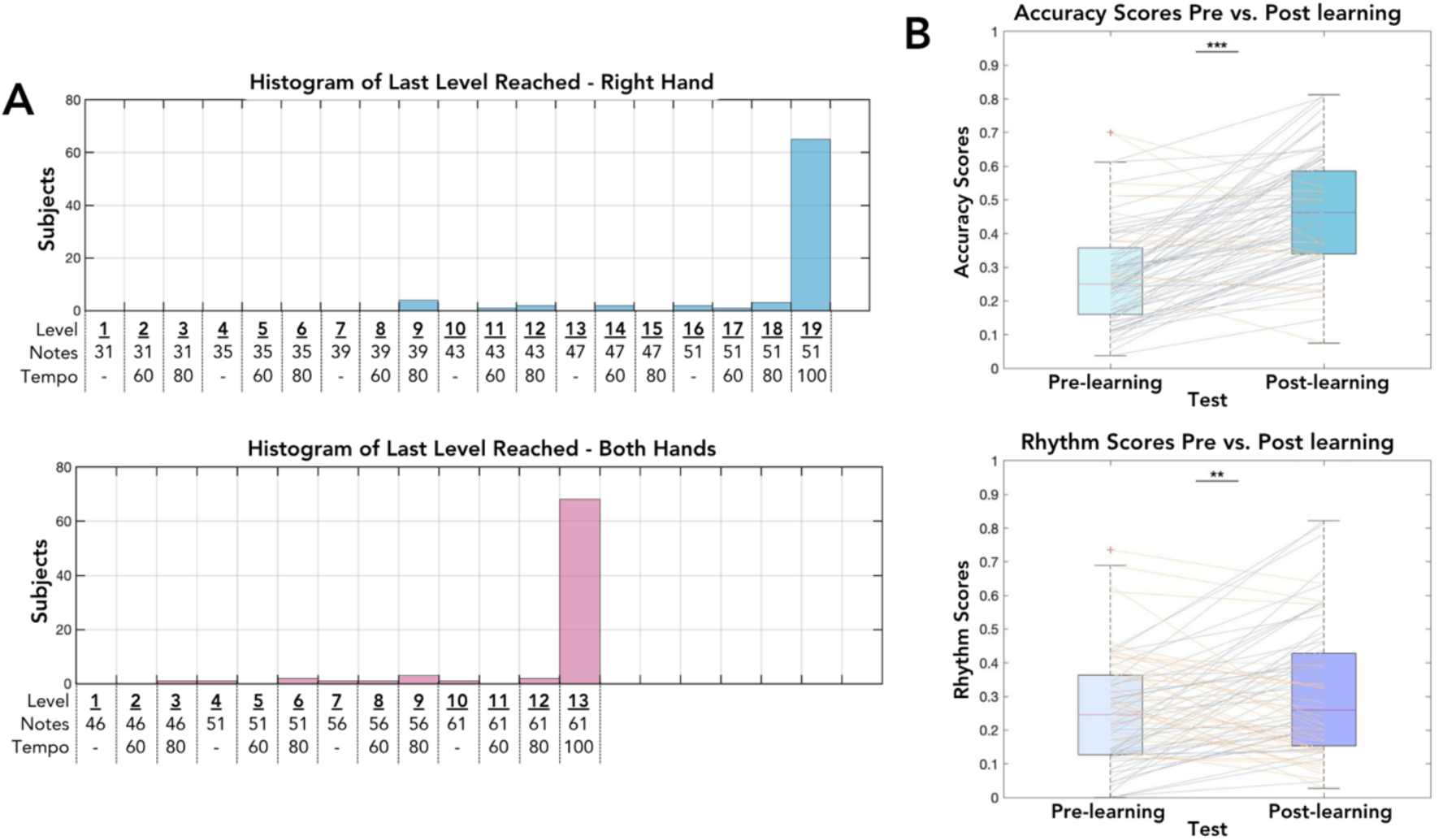
Behavioral improvements in learned and unlearned melodies. **A.** Histograms showing training levels completed by participants. The X axis describes the level number, the number of notes, and the tempo required at each level (‘-‘ represents a level without tempo requirements); the Y axis represents the number of participants who successfully completed each level of learning with the right hand only (Top) or with both hands (Bottom). **B.** Improvement in accuracy (Top) and rhythm scores (Bottom) in the generalization tests following learning. Each line represents an individual participant; gray lines indicate participants with improved performance following training, whereas orange lines represent participants with decreased performance. Red crosses represent outliers.

In addition to improved performance on the learned melody, participants also showed enhanced performance on the generalization test, which required playing an unfamiliar melody, as reflected by improvements in both rhythmic precision and accuracy scores (Figure 3B; Accuracy: Rhythm: ). No significant change was observed in tempo scores following learning ().

### Context-dependent brain activity changes following musical training

Following musical training, participants demonstrated increased brain activations in response to passive listening to *Für Elise* played on the piano, i.e., as in the training procedure. Brain activity to the “*Für Elise* > *Ode to Joy*” task-contrast, which includes both piano and saxophone presentations of the two melodies, also increased, suggesting changes at the melodic level, while no significant effects were detected at the instrumental level (*Für Elise* Piano > *Für Elise* Saxophone) (Table 1). This indicates that training-induced activity changes were mostly driven by sensitivity to the melodic content of the learned musical piece, rather than by the specific instrumental timbre (Figure 4).

**Figure 4.**
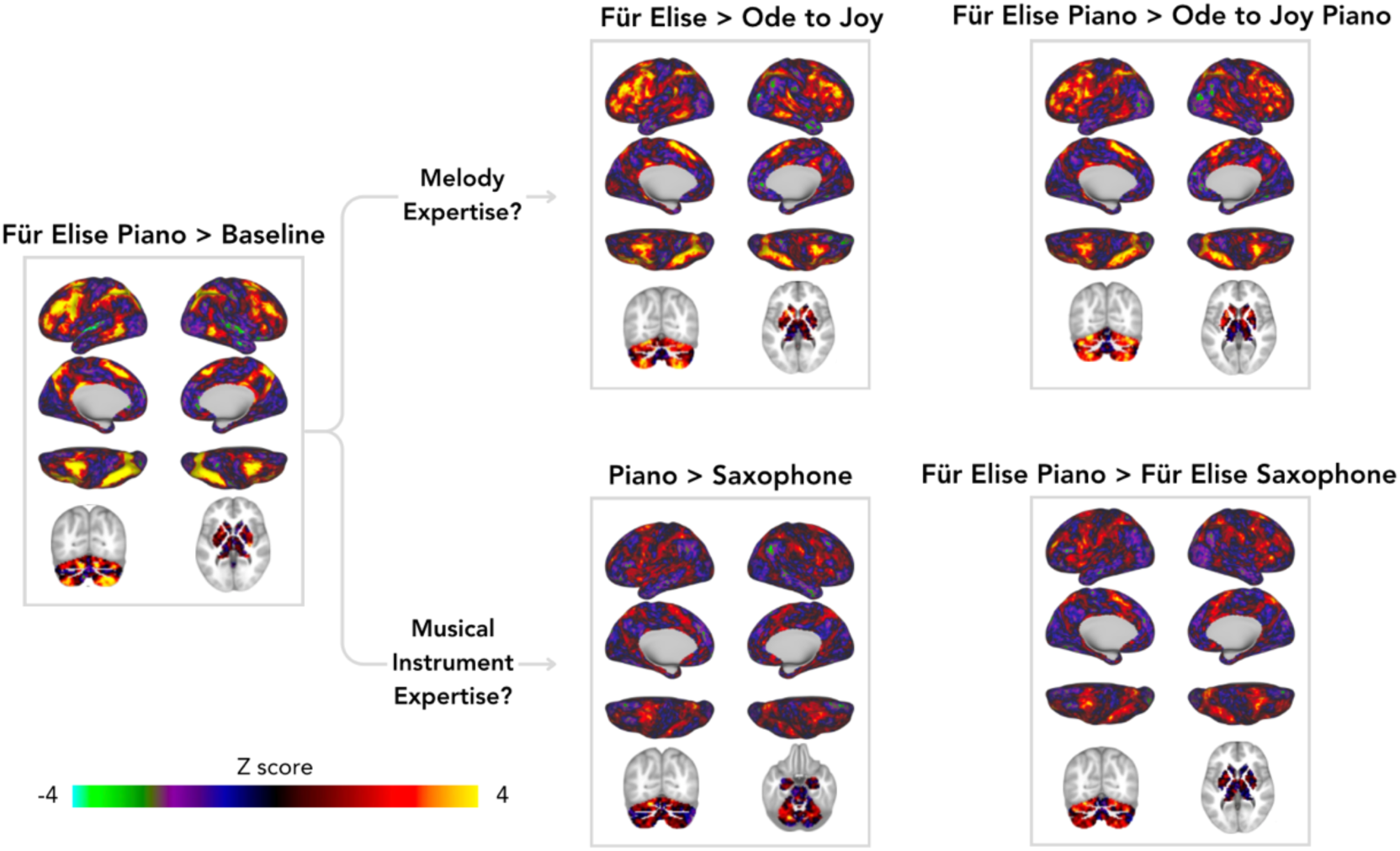
Context dependent changes in brain activity following musical training. Unthresholded Z-score maps showing learning-induced changes in brain activity for the different task-contrasts. Warm colors represent increased activity following learning; cool colors represent decreased activity. Overall, these findings suggest that activity changes were driven more by expertise with the specific melody rather than with the musical instrument.

**Table 1.**
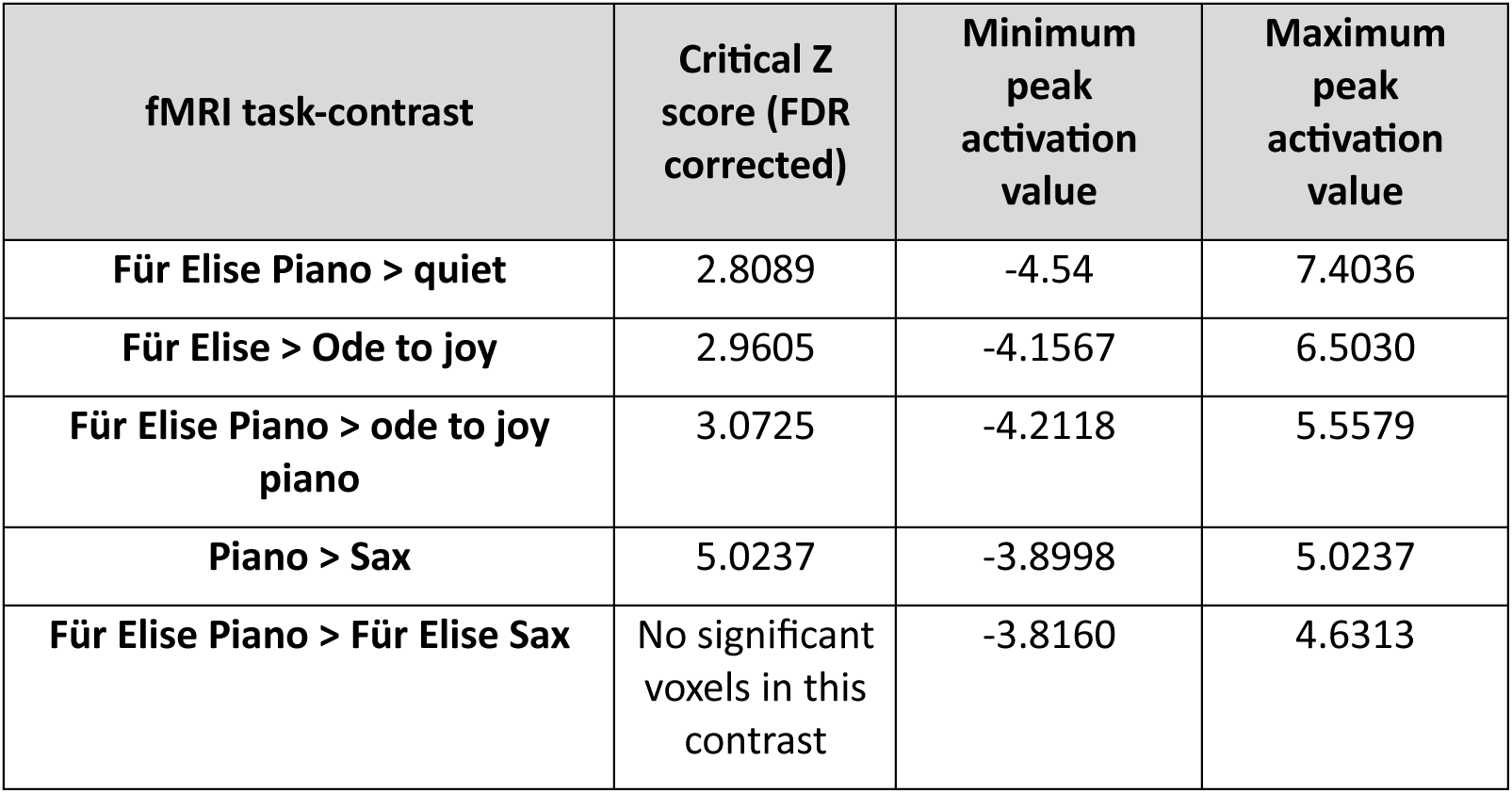
Whole-brain fMRI activation changes following musical training. Task-contrasts are ordered by the magnitude of activity change, from highest to lowest.

The spatial pattern of training-related neural changes was generally similar across contrasts, with overlapping regions mainly differing in effect size. Increases in activity were observed in skill-related areas such as the bilateral cerebellum, left intraparietal sulcus, and bilateral supplementary motor area, while decreases were noted in the bilateral superior temporal gyrus only in the learned context (*Für Elise* played on the piano > quiet). No changes in brain activity were observed in the passive control group.

### Classification of pre- vs. post-training brains

Classification models were trained on either the training or the control pre-learning fMRI data. We then tested whether pre- and post-learning scans can be differentiated based on whole- brain activation patterns. All models of the piano training group performed better than chance level, whereas models based on the passive control group did not exceed chance level in any of the task contrasts. Among the classification models of the training group, the model based on brain activity to the contrast “ *Für Elise* piano > quiet” yielded the highest AUC of 0.82 () and accuracy of 76%, demonstrating strong discriminative power between pre- and post-learning scans (Figure 5). The model based on “*Für Elise* saxophone > quiet” also performed well, with an AUC of 0.80 () and an accuracy of 73%. In comparison, the models trained on brain activity to “*Ode to Joy* piano > quiet” or “*Ode to Joy* saxophone > quiet” showed lower AUC values of 0.7 and 0.77, respectively, and accuracy levels of 63% and 69%, respectively, indicating more modest classification performance with near-significance AUC rates. Overall, models based on *Für Elise* played on the piano or saxophone, as well as the combined model, performed significantly better than models based on *Ode to Joy* (). These results suggest that whole-brain activity patterns, particularly in response to the learned musical piece, provide reliable indicators of learning-related neural changes.

**Figure 5.**
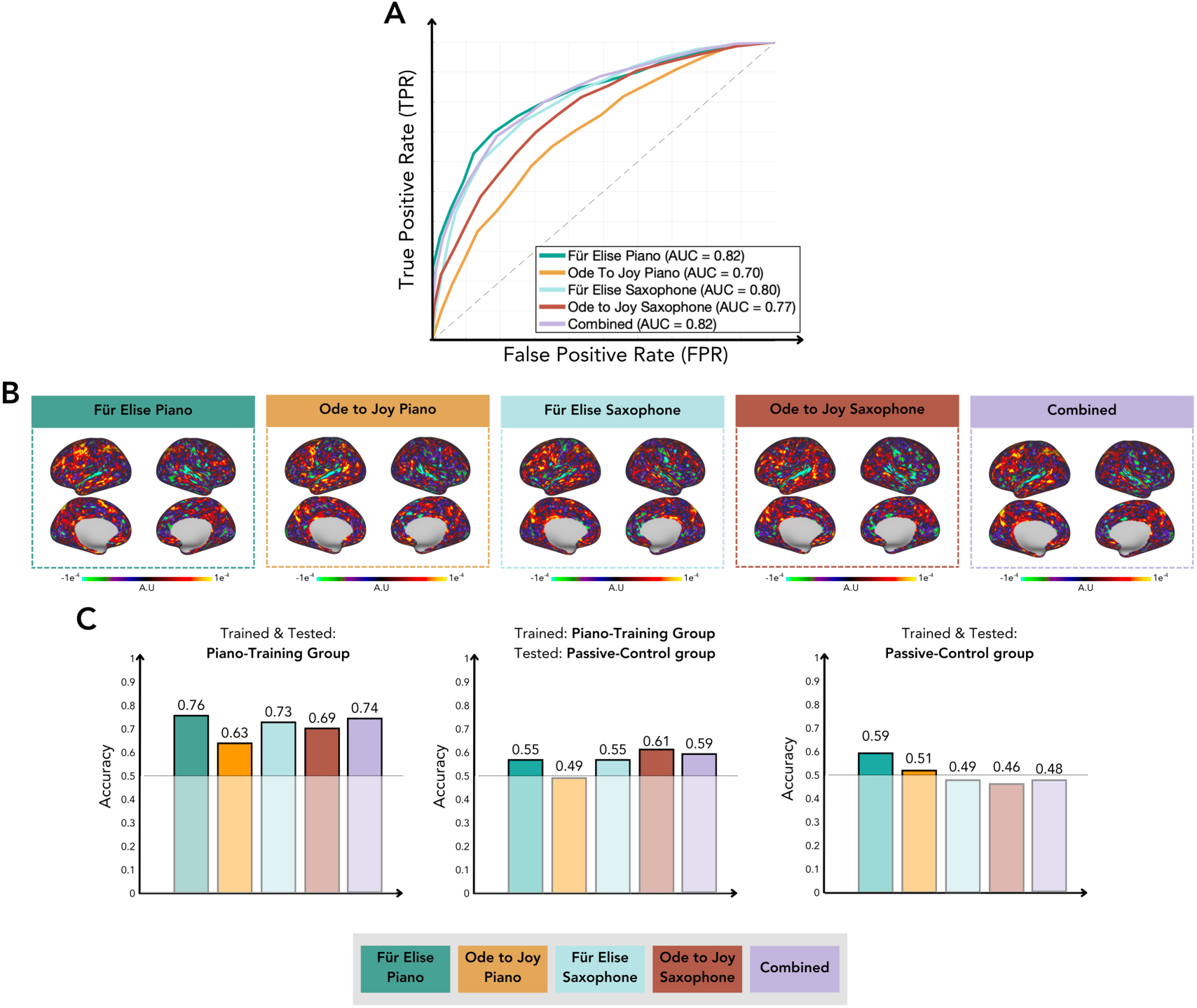
Classification of pre- vs. post-training scans. **A.** Classification performance evaluated on data from the piano-training group, using Receiver Operating Characteristic (ROC) Area Under the Curve (AUC) for models classifying pre- vs. post-training scans based on whole-brain activity patterns to the different task conditions contrasted against the quiet condition. The diagonal line indicates a random classifier’s performance (AUC = 0.5). Each curve represents mean performance over 100 cross-validation models (10 folds × 10 repetitions), using logistic regression with PCA. **B.** Contribution maps depicting brain areas that contributed the most to each model’s decision. Warmer colors indicate vertices with greater influence on models’ decision. **C.** Classification accuracy under different training- testing configurations: models trained and tested on the piano-training group (left), models trained on the piano-training group but tested on the passive control group (middle), and models trained and tested on the passive control group (right).

Pairwise comparisons revealed that the model based on “Für Elise piano > quiet” achieved significantly higher classification accuracy than the models based on “Ode to Joy piano > quiet” and “Ode to Joy saxophone > quiet” (). The “Für Elise saxophone > quiet” model also significantly outperformed both Ode to Joy models (). Interestingly, within the untrained melody conditions, the model based on “Ode to Joy saxophone > quiet” significantly outperformed the model based on “Ode to Joy piano > quiet” (69% vs. 63%, ). Finally, the combined model achieved significantly higher accuracy than both Ode to Joy models ().

### Pre-training brain activity predicts subsequent behavioral performance

We examined whether pre-training brain activity in response to the learned melody and instrument (i.e., *Für Elise* played on the piano) predicts behavioral performance following training. A significant positive correlation was found between actual and predicted accuracy scores in the final lesson (; Figure 6), indicating that pre-training neural responses shape subsequent performance. Predictions based on activation patterns to *Für Elise* played on the saxophone (), or to *Ode to Joy* played on the piano () or saxophone (), or the combined model () did not reach statistical significance. Predictions of rhythm scores based on pre-training brain activity to these task-contrasts were also not significant (all p > 0.05).

**Figure 6.**
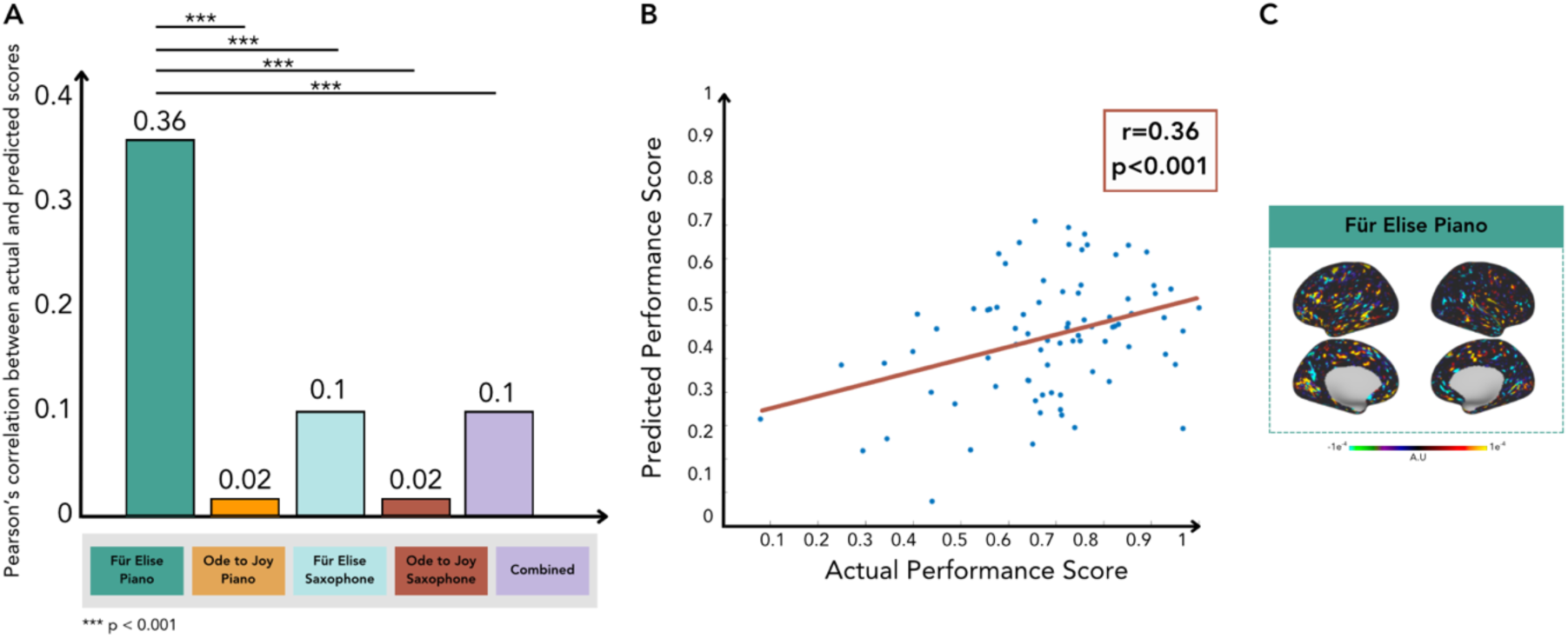
Pre-training activity patterns predict subsequent behavioral performance. **A.** Correlations between actual and predicted scores in the post-training generalization test. Prediction models were trained on pre-training brain activity to the different task conditions contrasted with quiet. **B.** Accuracy scores derived from pre-training brain activity patterns in response to *Für Elise* played on the piano significantly correlated with the actual scores. Each dot represents an individual participant; the red line represents the best-fit linear regression. **C.** Brain map showing areas that contributed the most to prediction of the post-training generalization test accuracy score from pre-training activity to *Für Elise* played on the piano. Warmer colors indicate vertices with greater influence on the model’s decision.

We next examined whether pre-training brain activity in response to musical instruments regardless of melody predicts post-training performance. To this end, we trained a prediction model on the Piano vs. Saxophone task-contrast, which captures the sensitivity to musical instruments prior to any piano training. This model achieved predicted rhythm scores which significantly correlated with the actual ones (; Figure 7), meaning that pre-training neural responses elicited by untrained musical instruments were predictive of rhythm performance in novel musical contexts after piano training. Predictions of accuracy scores from brain activity to the Piano vs. Saxophone task-contrast were not significant (p > 0.05).

**Figure 7.**
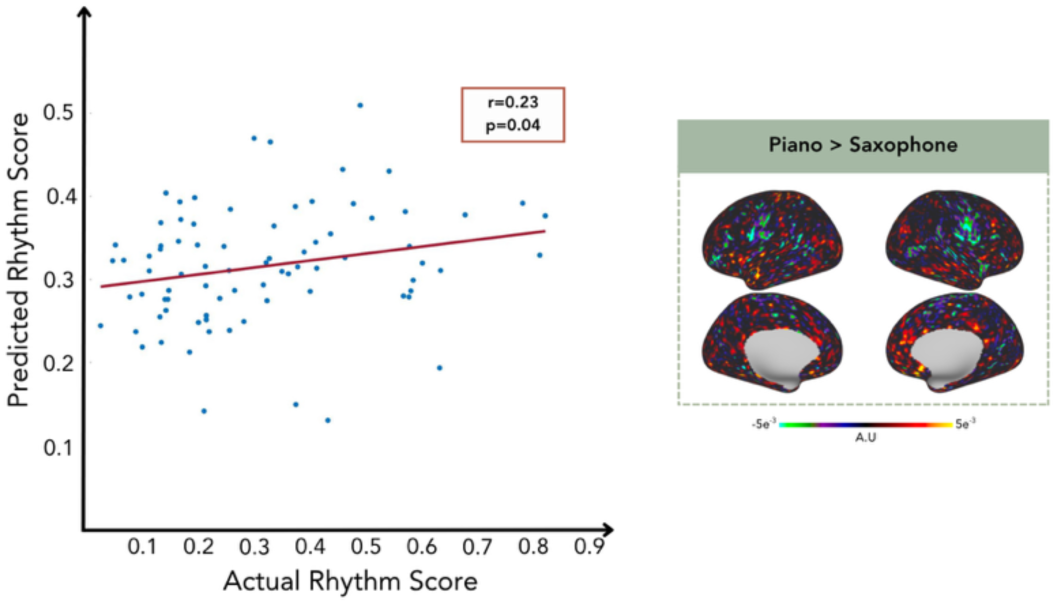
Pre-training sensitivity to musical instruments predicts post-training performance. Rhythm scores derived from pre- training brain activity patterns in response to the Piano > Saxophone task-contrast significantly correlated with the actual rhythm scores. Each dot represents an individual participant; the red line indicates the best-fit linear regression. Brain map showing areas that contributed the most to prediction; warmer colors indicate vertices with greater influence on the model’s decision.

### Classification of melodies and musical instruments improves following training

We tested whether whole-brain activity patterns pre- and post-training could be used to classify between melodies (*Für Elise* vs. *Ode to Joy*) and musical instruments (Piano vs. Saxophone) (Figure 8). A classifier trained on pre-learning brain activity of *Für Elise* played on piano vs. quiet and *Ode to Joy* played on piano vs. quiet achieved a mean AUC of 0.66 and accuracy of 0.65 for classification of *Für Elise* vs. *Ode to Joy.* A permutation test with 10,000 label shuffles indicated a non-significant decoding of melody prior to learning (). Post-training performance showed an AUC of 0.74 () and accuracy of 0.72, reflecting significantly higher neural separability between the two melodies ().

**Figure 8.**
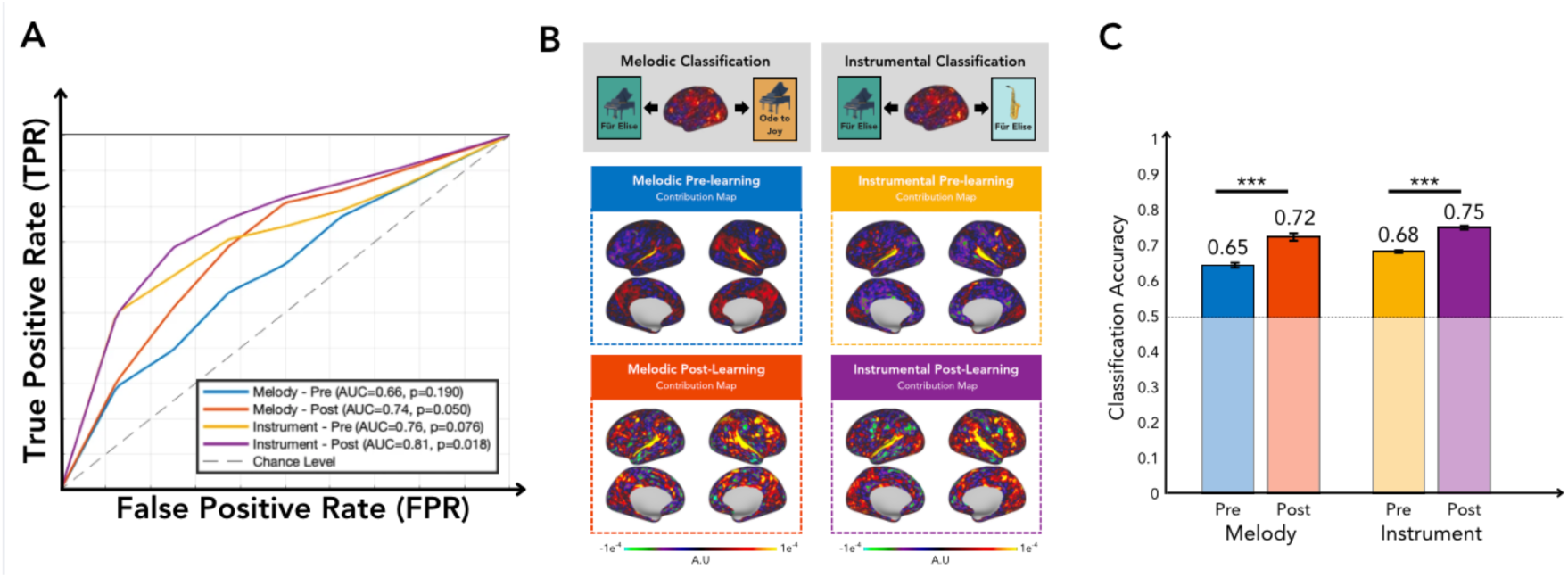
Classification of melodies and musical instruments. **A.** Receiver Operating Characteristic (ROC) curves show mean classifier performance for decoding of melody (*Für Elise* vs. *Ode to Joy*) and instrument (Piano vs. Saxophone) from whole-brain fMRI activation patterns before or after piano training. All models were trained and tested on the data from the piano-training group using repeated cross-validation (10 folds × 10 repetitions; 100 models). The dashed diagonal indicates chance-level performance. **B.** Contribution maps for the melodic and instrumental classification pre- and post-learning. Warmer colors indicate vertices with greater influence on the classifier’s decision. **C.** Mean classification accuracy (± SEM) of models trained and tested on the data from the piano-training group, before and after learning. The horizontal gray line indicates chance-level performance (0.5). Asterisks denote significant differences between classification performance before and after piano training ), assessed using pairwise t-tests across all folds and repetitions

For instrument classification, which used the *Für Elise* played on piano vs quiet and *Für Elise* played on saxophone vs quiet, the pre-training model yielded a mean AUC of 0.76 and accuracy of 0.68, suggesting non-significant decoding accuracy (). Classification based on post- training brain activity achieved an AUC of 0.81 () and accuracy of 0.75, indicating a statistically significant enhancement in decoding ability (t(99) = −5.09, p < .001).

Performance of models trained on data from the piano-training group and applied to the passive-control group was similar in the first time point but was consistently lower in the second time point, indicating no evidence of generalization of training-derived features to untrained participants. In this external test, melody classification accuracy remained stable across sessions (0.59 pre- and post-training) and instrument classification showed a non- significant decrease (0.69 pre-training to 0.62 post-training).

## Discussion

This study examines two fundamental aspects of musical skill acquisition: whether it is context-specific or generalized beyond the trained melody and instrument, and whether pre- training brain activity is predictive of subsequent performance. Previous studies have employed a range of experimental approaches, from highly controlled motor learning paradigms to more naturalistic musical training protocols. In the present study, we combined a complex, real-world musical task with dense behavioral tracking. Our findings support a melodic rather than instrumental expertise in musical training, as reflected by (1) training- induced activity changes that are mostly specific to the learned musical content and (2) classification between pre- and post-training brains that is mostly driven by activity patterns in response to the learned musical content. Predictions of musical performance following training from pre-training brain activity revealed a more complex picture, such that prediction of accuracy is relatively melody-specific while prediction of rhythm is more broadly based on sensitivity to musical instruments.

Following training, activity in the superior temporal gyrus during passive listening to *Für Elise* played on the piano (i.e., the trained melody and instrument) significantly decreased. This finding is consistent with the idea of neural efficiency, according to which the processing of a familiar (trained) stimulus may require fewer resources compared to a novel (untrained) stimulus (Chen et al., 2012). In converse, increased activations following training were observed in the bilateral cerebellum, left intraparietal sulcus, and bilateral supplementary motor area, reflecting the strengthening of auditory-motor integration (Brown and Penhune, 2018; Li et al., 2018; Olszewska et al., 2021; Zatorre et al., 2007).

Differential neural responses to familiar and unfamiliar auditory stimuli have been repeatedly demonstrated in fMRI (Freitas et al., 2018) as well as EGG and MEG studies (Belo et al., 2023; Kobayashi et al., 2024), indicating higher audio-motor synchronization for familiar tunes. Consistent with the findings reported here, listening to trained compared to untrained melodies has been shown to elicit increased activations in the bilateral premotor and dorsolateral prefrontal cortices, the posterior parietal cortex, and the cerebellum (Olszewska et al., 2021). The dorsolateral prefrontal cortex, together with parietal regions, has been suggested to store representations of learned sequences (Doyon and Benali, 2005; Penhune and Steele, 2012), also reflecting the engagement of the reward system while listening to familiar music (Fasano et al., 2023). Since the process of musical skill acquisition inherently involves the transition of a trained musical piece from unfamiliar (or less familiar) to familiar, the training-induced changes in the response to *Für Elise* played on the piano were expected. Importantly, however, training-induced changes were less pronounced in response to the trained instrument alone, suggesting that musical expertise may be more closely tied to the specific temporal structure of melody than to the instrumental characteristics.

A notable exception was observed for the untrained melody: pre- versus post-training classification was more accurate when Ode to Joy was played on the saxophone than on the piano. One possible explanation is that pairing the untrained melody with the training- relevant piano timbre caused varying levels of generalization among participants. As a result, post-training representations might have shifted in some participants while remaining closer to the pre-training patterns in other, thus increasing inter-subject variability. This interpretation is consistent with evidence that familiar instrumental timbres can activate prior musical knowledge and bias the processing of novel tonal sequences (Van Hedger et al., 2022). The decreased classifier performance for Ode to Joy played on the piano may therefore be explained by an increased overlap between pre- and post-training patterns at the group level.

While the neural reorganization induced by training seems to be largely driven by the acquisition of specific melodic sequences rather than a generalized enhancement in processing the piano’s timbre, we did observe generalization of the acquired skill at the behavioral level. Specifically, participants improved their performance of an unfamiliar melody in the post-training generalization test, indicating that behavioral gain was not limited to *Für Elise*. This dual pattern of melodic specificity in neural modifications on one hand, and generalization in behavioral performance on the other, suggests that while our brain may encode a specific learned content, the underlying motor and rhythmic skills that are being acquired may be transferable to novel contexts (Aagten-Murphy et al., 2014; Rohrmeier and Rebuschat, 2012; Van Hedger et al., 2023). For example, it has been recently shown that pitch judgments of a familiar melody are relatively robust across varying timbre and octave (Van Hedger et al., 2023). There is also, however, conflicting evidence showing limited generalizability of performance under inverted pitch conditions (Pfordresher et al., 2021), warranting further research on the degree to which musical training generalizes at the behavioral level. On a broader perspective, the acquisition of musical skills has been linked to behavioral improvements across auditory, spatial and linguistic domains (Wang, 2022) suggesting potential transfers of the acquired abilities beyond musical performance per se (Rymarczyk and Cybulska, 2025; Smit et al., 2023).

Our results provide evidence for a neural predisposition to musical learning. Pre-training brain activity in response to the trained melody, as measured *before training*, predicted subsequent playing accuracy, while sensitivity to instrumental timbre (Piano vs. Saxophone) before training predicted rhythmic performance. This distinction suggests that different aspects of musical learning may rely on complementary representations: Whereas melody-specific representations appear to primarily support the acquisition of expertise for the practiced material, broader auditory representations may contribute to successful learning and facilitate behavioral transfer beyond the trained melody. These findings suggest that the initial state of the brain should not be viewed merely as a baseline but as a core element that shapes learning trajectory and influences its outcomes (Herholz et al., 2016; Olszewska et al., 2021; Wollman et al., 2018; Zuk and Gaab, 2018). Predispositions for musical training have so far been scarcely studied, with more emphasis put on behavioral tendencies (e.g., perceived competence, mindset and aptitude; Bianco et al., 2019) or general intelligence (Burgoyne et al., 2019) than on neural markers. Our findings underscore the importance of individual differences in pre-existing activation patterns for predicting skill acquisition. Future studies may address the utility of other neural properties, including brain structure and connectivity, as predictors of musical learning abilities (Zatorre, 2013).

Prediction of musical performance from pre-training neural properties may reflect the quality of perceptual encoding prior to training, such that individuals who exhibit greater sensitivity to relevant auditory features form more precise neural representations that support subsequent learning (Herholz and Zatorre, 2012; Zatorre, 2013). Specifically, prediction of accuracy appears to rely more on sensitivity to sequence-related features, such as pitch discrimination and the structured organization of melodic content, consistent with evidence that stable melodic representations and enhanced pitch processing support musical expertise (Bianchi et al., 2017; Van Hedger et al., 2023). In contrast, predicting rhythm is better explained by sensitivity to more general acoustic features, such as onset characteristics and the temporal envelope, which govern how sounds emerge and decay over time and are closely linked to the extraction of temporal regularities in auditory-motor timing networks (Toiviainen et al., 2014; Vuust et al., 2022). These features are essential for accurate timing and indicate that attentional sensitivity to sound dynamics, rather than to its melodic structure alone, significantly influences rhythmic performance.

Finally, we demonstrate that training enhances decoding of melodic and instrumental information from neural patterns. Specifically, classification models showed significantly higher neural separability between different melodies and instruments following training. This suggests that musical practice not only changes the magnitude of brain responses but also refines the distinctiveness of neural representations for musical features. These results are in line with previous studies that showed successful decoding of musical pieces (Hoefle et al., 2018) and instruments (Cheung et al., 2023) from brain activity patterns. Also consistent with previous reports, brain areas that contributed most to classification from pre-training data included the bilateral superior temporal gyrus and Heschl’s gyrus (Toiviainen et al., 2014), while following learning additional motor and prefrontal regions came into play. The contribution of motor regions to classification, especially that of musical instruments, again emphasizes the strengthening of auditory-motor integration following musical training (Herholz et al., 2016; Olszewska et al., 2021; Zatorre et al., 2007).

The gain in classification of melodies following training complements our main finding, that musical training reflects expertise in a practiced melody more than the abstract rules of a musical instrument. Nevertheless, the gain in classification of instruments suggests that training also induced some degree of instrumental characteristics proficiency, at least to the extent that enhanced classification performance post-training.

## Conclusions

We conclude that pre-existing neural characteristics interact with practice-induced functional neuroplasticity to shape musical skill acquisition. Our findings suggest that practice-induced plasticity predominantly refines melody-specific neural representations, whereas broader pre-existing neural characteristics may facilitate successful learning and behavioral generalization. Our results indicate that while behavioral skills acquired during musical training can generalize to novel contexts, the neural modifications induced by training are predominantly specific to the learned melodic content. Future research should explore whether longer training periods or more diverse repertoires eventually lead to a shift from melodic-specific to instrument-generalized neural expertise.

